# Mitochondrial DNA variation of Leber’s Hereditary Optic Neuropathy (LHON) in Western Siberia

**DOI:** 10.1101/744219

**Authors:** E.B. Starikovskaya, S.A. Shalaurova, S.V. Dryomov, A.M. Nazhmidenova, N.V. Volodko, I.Y. Bychkov, I.O. Mazunin, R.I. Sukernik

## Abstract

Leber’s hereditary optic neuropathy (LHON) is a form of disorder caused by pathogenic mutations in a mitochondrial DNA. LHON is maternally inherited disease, which manifests mainly in young adults, affecting predominantly males. Clinically LHON has a manifestation as painless central vision loss, resulting in early onset of disability. Epidemiology of LHON has not been fully investigated yet. In this study, we report 44 genetically unrelated families with LHON manifestation. We performed whole mtDNA genome sequencing and provided genealogical and molecular genetic data on mutations and haplogroup background of LHON patients in the Western Siberia population. Known “primary” pathogenic mtDNA mutations (MITOMAP) were found in 32 families: m.11778G>A represents 53,10% (17/32), m.3460G>A – 21,90% (7/32), m.14484T>C – 18,75% (6/32), and rare m.10663T>C and m.3635G>A represent 6,25% (2/32). We describe potentially pathogenic m.4659G>A in one subject without known pathogenic mutations, and potentially pathogenic m.9444C>T, m.6261G>A, m.9921G>A, m.8551T>C, m.8412T>C, m.15077G>A in families with known pathogenic mutations confirmed. We suppose these mutations could contribute to the pathogenesis of optic neuropathy development. Our results indicate that haplogroup affiliation and mutational spectrum of the Western Siberian LHON cohort substantially deviate from those of European populations.

## Introduction

Leber’s hereditary optic neuropathy is a form of hereditary disorder caused by pathogenic mutations in a mitochondrial DNA. These mutations are non-synonymous and affect genes coding for different subunits of complex I of the mitochondrial respiratory chain. The occurrence of such kind of mutations in mtDNA subunits leads to dysfunction of the electron transport, increased reactive oxygen species production and defective ATP synthesis (Baracca, Solaini et al. 2005; Lin, Sharpley et al. 2012; Kim, Jurcute and Yu-Wai-Man 2018). Retinal ganglion cells are highly susceptible to death during LHON progression, because of their high sensibility to disrupted ATP production and oxidative stress (Meyerson, Van Stavern et al. 2015). Therefore, LHON is usually manifested as painless, acute or subacute, with central visual loss in both or one eye, results to early onset of disability. The onset of Leber hereditary optic neuropathy is relatively rare in childhood and has better visual prognosis. The peak age of onset of visual loss among LHON carriers is 20–30 years old (Majander, Bowman et al. 2016). In some cases, LHON patients have been reported as having additional neurologic, cardiac, and endocrine disorders (Finsterer and Zarrouk-Mahjoub 2016; Haas 2019, de Barcelos, Troxell and Graves 2019). Leber’s disease is inherited maternally and manifests in young adults, predominantly affecting males – 50% of male and only 10% of female subjects experience vision loss. Interestingly, this sex predilection cannot be explained by the principles of mitochondrial inheritance (Meyerson, Van Stavern et al. 2015). At the present time a number of mtDNA point mutations have been described, but the most prevalent are m.3460G>A, m.11778G>A and m.14484T>C, accounting for about 90% of cases of LHON worldwide. The prevalence of each mutation varies among different populations, but the average is 69-92% for m.11778G>A, 3-19% for m.14484T>C and 1-13% for m.3460G>A (Mackey, Oostra et al. 1996; Mashima, Yamada et al. 1998; Kumar, Kaur et al. 2012, Romero, Fernandez et al. 2014; Jiang, Liang et al. 2015; Khan, Govindaraj et al. 2017). However, there are significant deviations from the average in some populations, for example, among French Canadians 87% of cases are due to m.14484T>C as a result of a founder effect (Laberge, Jomphe et al. 2005). Moreover, phenotypic expression of these primary mutations has been found to vary in different populations and different pedigrees. This incomplete penetrance suggests that other factors, such as mtDNA haplogroup background, nuclear genetic background and environmental factors, may influence the modulation of phenotypic expression and severity of the disease (Jacobi, Leo-Kottler et al. 2001; Kirkman, Yu-Wai-Man et al. 2009; Istikharah, Tun et al. 2013).

Consequently, the worldwide prevalence of LHON varies in different populations and is unknown for the majority of them. According to broad epidemiologic studies, it is estimated between 1:30000 and 1:50000 (Puomila, Hamalainen et al. 2007; Mascialino, Leinonen et al. 2012). Epidemiology of LHON has not been fully investigated in Russian Federation, and our previous studies included the description of isolated cases (Brown, Zhadanov et al. 2001; Brown, Starikovskaya et al. 2002; Volod’ko, L’vova et al. 2006). Hence, in the present study, we reported 44 genetically unrelated LHON families, performed whole mtDNA sequencing and provided genealogic and molecular genetic data on mutations and haplogroup background of LHON patients in the Western Siberia population.

## Materials and methods

### Subjects

This study was approved by the Ethics Committee IRB 00001360 affiliated with Vector State Research Center of Virology and Biotechnology (SRC VB Vector), Novosibirsk, Russian Federation. The total number of subjects in the study is 178 individuals from 44 unrelated families (85 affected and 83 healthy carriers), including 17 cases from our previous studies (Brown, Zhadanov et al. 2001; Brown, Starikovskaya et al. 2002; Volod’ko, L’vova et al. 2006). The clinical follow-up of LHON patients has been carried out by the Novosibirsk Branch of Federal Eye Microsurgery Department since 1997, conducted by one of the authors. The clinical diagnosis was based on a combination of symptoms and signs: painless acute or subacute central vision loss, fundus changes and visual field abnormality such as pseudopapilledema, optic nerve atrophy, central or cecocentral scotoma. All the individuals made an informed decision to take part in the study and provided written consent. Family history was taken in each case to identify maternal inheritance of symptoms. The complete mtDNA genomes were sequenced for the family’s probands, and for the other individuals the certain mutations were confirmed by sequencing of associated mtDNA regions.

### mtDNA analysis

Whole peripheral blood samples were collected from the donors in 10-ml Vacutech EDTA tubes. Total DNA was extracted from buffy-coat layer using the SileksMagNA-G(tm) Blood DNA Isolation kit, according to the manufacturer’s protocols. The complete sequencing procedure entailed PCR amplification of 22 overlapping mtDNA templates (Nochez, Arsene et al. 2009), which were sequenced in both directions with BigDye 3.1 terminator chemistry (PE Applied Biosystems, Foster City, CA, USA). The trace files were analyzed with Sequencher (version 4.5 GeneCode Corporation) software. To perform capillary electrophoresis on ABI Prism 3130XL DNA Analyzer we used core facilities of the “Genomika” Sequencing Center, SBRAS, Novosibirsk, Russian Federation. Variants were scored relatively to the Reconstructed Sapiens Reference Sequence, RSRS (Behar, van Oven et al. 2012). MtDNA haplotypes were identified following the nomenclature suggested by the PhyloTree Build 17 (van Oven and Kayser 2009). 42 mitochondrial genomes obtained through this study were deposited in GenBank with accession numbers XX000000-XX000000. Two genomes, EU807741.1 and EU807742.1, had been accessed earlier (Brown, Zhadanov et al. 2001).

### Penetrance analysis

We determined penetrance as the proportion of affected individuals from all maternally related family members using family pedigrees (Puomila, Hamalainen et al. 2007). Values for both men and women were calculated separately. Pedigrees are available in supplementary materials.

### Analysis of pathogenicity for non-synonymous mutations

To make sure that the revealed non-synonymous mtDNA mutations are not sequencing errors or hot points for the general population, and to find out the disease–associated polymorphisms among those published earlier, we used several databases: MITOMAP (Brandon, Lott et al. 2005), mtDB - Human Mitochondrial Genome Database, containing 2704 human mitochondrial genomes (Ingman and Gyllensten 2006) and HmtDB – Human Mitochondrial DataBase, containing 32922 human mitochondrial genomes (Attimonelli, Accetturo et al. 2005). To assess the possible pathogenicity of these mutations and to predict whether a protein sequence variation affects protein function, we used the following web applications: MutPred 1.2 (Li, Krishnan et al. 2009), MutPred 2 (Pejaver et al. 2017), PolyPhen – 2 (Adzhubei, Schmidt et al. 2010), PROVEAN (Protein Variation Effect Analyzer) (Choi, Sims et al. 2012) and SIFT (Sorting Intolerant from Tolerant) (Kumar, Henikoff et al. 2009) programs. All the sources are provided in the public domain.

## Results

From 44 LHON families, 32 harbored a primary mutation; the results are shown in Table 1. Among families with a primary mutation, the m.11778G>A represents 53,10% (17/32), m.3460G>A – 21,90% (7/32), m.14484T>C – 18,75% (6/32). Rare m.10663T>C and m.3635G>A represent 6,25% (2/32) of the families.

**Table 1.** Summary of the data for examined LHON families. Published previously by: * (Volod’ko, L’vova M et al. 2006); ** (Brown, Starikovskaya et al. 2002; Volod’ko, L’vova et al. 2006); *** (Brown, Zhadanov et al. 2001; Volod’ko, L’vova et al. 2006).

| No. | Family name | Ethnicity on maternal line | Number of examined individuals (affected/ healthy) | Family history of visual loss | Primary mutation | mtDNA haplogroup |
| --- | --- | --- | --- | --- | --- | --- |
| 1 | L1* | Russian | 9 (1/8) | Yes | m.11778G>A | T2b |
| 2 | L2** | -/- | 6 (2/4) | Yes | m.10663T>C | J1c4 |
| 3 | L3* | -/- | 4 (1/3) | Yes | m.11778G>A | T2d1b1 |
| 4 | L5* | -/- | 1 (1/0) | Yes | m.11778G>A | J1c2i |
| 5 | L6 | -/- | 2 (2/0) | No | - | U4a1d |
| 6 | L8 | Unknown | 1 (1/0) | No | - | U2e1 |
| 7 | L9 | Russian | 2 (2/0) | Yes | - | U5a1b1c1 |
| 8 | L10* | -/- | 8 (3/5) | Yes | m.14484T>C | M9a1a1c1a |
| 9 | L12* | -/- | 2 (1/1) | No | m.11778G>A | J2b1a1 |
| 10 | L14* | -/- | 5 (2/3) | No | m.11778G>A | T2b28 |
| 11 | L17* | -/- | 8 (4/4) | Yes | m.14484T>C | J1c2c1 |
| 12 | L18* | Altaiian | 4 (2/2) | No | m.3460G>A | D4p |
| 13 | L20 | -/- | 3 (2/1) | Yes | - | U4b1b1 |
| 14 | L23* | Azerbaijani | 1 (1/0) | Unknown | m.11778G>A | J2b1 |
| 15 | L24* | Tuviniian | 19 (7/12) | Yes | m.3460G>A | C5d1 |
| 16 | L25*** | Russian | 5 (3/2) | Yes | m.3460G>A | D5a2a2 |
| 17 | L26* | -/- | 13 (8/5) | Yes | m.11778G>A | J1c7a |
| 18 | L27*** | -/- | 11 (2/9) | Yes | m.11778G>A | H2a5b |
| 19 | L28*** | -/- | 9 (2/7) | Yes | m.11778G>A | T2b8 |
| 20 | L30*** | -/- | 19 (11/8) | Yes | m.3635G>A | J2b1c1 |
| 21 | L31*** | -/- | 3 (2/1) | Yes | - | U3b1b |
| 22 | L32 | Unknown | 1 (1/0) | Unknown | m.14484T>C | V |
| 23 | L38 | Ukrainian | 2 (2/0) | Yes | m.11778G>A | J1c2c2a |
| 24 | L39 | Unknown | 2 (1/1) | No | m.11778G>A | V |
| 25 | L40 | Albanian | 1 (1/0) | Yes | m.14484T>C | H |
| 26 | L41 | German | 2 (1/1) | No | m.3460G>A | H40a |
| 27 | L42 | Belarusian | 2 (1/1) | Yes | m.11778G>A | H1c |
| 28 | L43 | Russian | 3 (1/2) | No | m.11778G>A | H |
| 29 | L45 | -/- | 2 (1/1) | No | - | U5a2e |
| 30 | L46 | -/- | 2 (1/1) | No | - | H13a1d |
| 31 | L47 | -/- | 1 (1/0) | No | m.14484T>C | J1c5a1 |
| 32 | L49 | Unknown | 1 (1/0) | No | m.11778G>A | K1c |
| 33 | L50 | Unknown | 1 (1/0) | Unknown | m.14484T>C | U5a2b1c |
| 34 | L51 | Unknown | 1 (1/0) | Unknown | - | U2c1b |
| 35 | L52 | Russian | 3 (1/2) | No | m.11778G>A | J1c2 |
| 36 | L53 | Unknown | 3 (1/2) | Unknown | m.11778G>A | H1b2 |
| 37 | L54 | Russian | 2 (1/1) | No | - | U4a2a |
| 38 | L56 | -/- | 2 (1/1) | No | - | J1c1b1 |
| 39 | L57 | -/- | 1 (1/0) | Yes | m.3460G>A | V1a1 |
| 40 | L58 | -/- | 2 (1/1) | Yes | m.3460G>A | J1c3 |
| 41 | L59 | Ukrainian | 2 (1/1) | No | - | V7a |
| 42 | L60 | Russian | 2 (1/1) | No | m.11778G>A | H1b2 |
| 43 | L61 | -/- | 4 (2/2) | Yes | m.3460G>A | H1b1 |
| 44 | L62 | -/- | 1 (1/0) | Yes | - | H |

According to the family pedigrees, only 50% (22/44) of cases had a family history of vision loss on maternal lineage in more than one generation, among which m.11778G>A represented 36% (8/22), the m.14484T>C – 14% (3/22), m.3460G>A – 23% (5/22) and without primary mutations (LHON-like cases) – 18% (4/22). Rare mutation cases (m.10663T>C and m.3635G>A) were family–inherited. 39% (17/44) of cases were sporadic, among which 13 cases were with only one affected person diagnosed and 4 cases with two affected persons in one generation. Among the sporadic cases m.11778G>A represents 41% (7/17), m.3460G>A – 12% (2/17), m.14484T>C – 6% (1/17), and the LHON-like cases – 41% (7/17).

Summary information on penetrance is shown in Table 2. The average penetrance among men was 32% (6-100%) and among women – 12% (0-58%). Our data correlate with data previously published by (Meyerson, Van Stavern et al. 2015). However, there are particular families with higher penetrance among females than among males: L24, L26, L28. The penetrance is highly variable between separate families, even with the same primary mutation as shown in pedigrees in supplementary materials.

**Table 2.** Summary information about penetrance. Pedigrees are available in supplementary materials.

|  | m.11778G>A (N=15) | m.14484T>C (N=4) | m.3460G>A (N=7) | Total |
| --- | --- | --- | --- | --- |
| Males | 34% | 46% | 15% | 31% |
| Females | 12% | 12% | 10% | 11% |

In 12 families with a clear-cut LHON phenotype, no pathogenic mtDNA mutations were found. Analysis of the mtDNA revealed non-synonymous mutations: m.14002A>G, m.4766A>G, and m.13105A>G, which have not been noted as associated with LHON or other diseases in the MITOMAP database. All of pathogenicity prediction tools indicated low probability that these amino acid substitutions are disease-associated for these mutations. Mutation m.4659G>A has been previously reported as associated with Parkinson’s disease (Khusnutdinova, Gilyazova et al. 2008), and in an Australian LHON pedigree that was heteroplasmic for the m.14484T>C (Mackey and Howell 1992). Polyphen-2 predicted the pathogenicity for this mutation as benign, MutPed 2 showed low probability score, but MutPred 1.2, PROVEAN and SIFT determined this mutation as deleterious. The results are shown in Table 3.

**Table 3.** Non-synonymous mutations revealed in LHON–like cases. Known primary mutations (m.11778G>A, m.3460G>A, m.14484T>C, m.10663T>C, m.3635G>A) were placed into this table to demonstrate differences between different prediction algorithms and frequencies in general population for pathogenic mutations. MutPred 1.2 score >0,75 referred to as confident hypotheses to be pathogenic. Mutpred 2 score >0.50 would suggest pathogenicity.

| Mutation | Protein-coding region of mtDNA | Amino acid substitution | PolyPhen – 2 score | MutPred 1.2/2 Score (cutoff 0,75/0,50) | PROVEAN/ SIFT prediction | Frequency in general population (mtDB) | Frequency in general population (HmtDB) | Family |
| --- | --- | --- | --- | --- | --- | --- | --- | --- |
| m.14002A>G | ND5 | T556A | 0,002 (benign) | 0,387/0,059 | Neutral/Tolerated | 0,0037 | 0,00289 | L45 |
| m.4766A>G | ND2 | M99I | 0,001 (benign) | 0,571/0,225 | Neutral/Tolerated | 0 | 0,00009 | L46 |
| m.13105A>G | ND5 | I257V | 0,001 (benign) | 0,198/0,032 | Neutral/Tolerated | 0,0612 | 0 | L51 |
| m.4659G>A | ND2 | A64T | 0,029 (benign) | 0,790/0,256 | Deleterious/Damaging | 0,0011 | 0,00161 |  |
| m.11778G>A | ND4 | R340H | 0,999 (probably damaging) | 0,919/0,494 | Deleterious/Damaging | 0,0097 | 0,00340 | - |
| m.3460G>A | ND1 | A52T | 1,000 (probably damaging) | 0,789/0,418 | Neutral/Damaging | 0,0097 | 0,00058 | - |
| m.14484T>C | ND6 | M64V | 0,993 (probably damaging) | 0,618/0,787 | Neutral/Damaging | 0,0026 | 0,00146 | - |
| m.10663T>C | ND4L | V65A | 0,946 (probably damaging) | 0,604/0,694 | Deleterious/Damaging | 0 | 0,00003 | - |
| m.3635G>A | ND1 | S110N | 0,999 (probably damaging) | 0,873/0,493 | Deleterious/Damaging | 0 | 0,00027 | - |

In several families with primary mutations (L01, L03, L12, L28, L30, L40, L43) we found out additional non-synonymous mutations, as shown in Table 4. Mutations m.8875T>C, m.14582A>G, m.8400T>C, m.4639T>C were neutral and mutation m.9444C>T had high probability to be pathogenic according to data of all the pathogenicity prediction tools; for other mutations we observed divergence of prediction results. Since prediction results for primary pathogenic mutations diverged too (see Table 3), novel non-synonymous nucleotide change was considered potentially pathogenic if it had extremely low frequency in the general population and it was predicted by at least three algorithms to have an effect on protein function. For mutations m.6261G>A and m.15468C>T only PolyPhen2 predicted pathogenicity as probably damaging and possible damaging respectively. However, mutation m.6261G>A has already been reported by Abu-Amero (Abu-Amero and Bosley 2006) in patient with optic neuropathy and also as somatic mutation associated with prostate cancer. Interestingly, that the family (L01) with m.6261G>A and m.11778G>A has the same haplogroup – T2, as case was reported by Abu-Amero. Other mutations (m.9921G>A, m.8551T>C, m.8412T>C, m.15077G>A) were predicted as pathogenic at least by three algorithms, but the first three of them have not been noted as associated with diseases in the MITOMAP database, and mutation m.15077G>A was reported as associated with maternally inherited isolated deafness (Gutierrez Cortes, Pertuiset et al. 2012).

**Table 4.** Additional non-synonymous mutations revealed in LHON cases. MutPred 1.2 score >0,75 referred to as confident hypotheses to be pathogenic. Mutpred 2 score >0.50 would suggest pathogenicity.

| Mutation | Protein-coding region of mtDNA | Amino acid substitution | PolyPhen – 2 score | MutPred 1.2/2 Score (cutoff 0,75/0,50) | PROVEAN /SIFT prediction | Frequency in general population (mtDB) | Frequency in general population (HmtDB) | Family |
| --- | --- | --- | --- | --- | --- | --- | --- | --- |
| m.6261G>A | CO1 | A120T | 0,998 (probably damaging) | 0,491/0,324 | Neutral/Tolerated | 0,0048 | 0,00553 | L01 |
| m.8875T>C | ATP6 | F117L | 0 (benign) | 0,251/0,429 | Neutral/Tolerated | 0,0007 | 0,001276 | L03 |
| m.9921G>A | CO3 | A239T | 0,009 (benign) | 0,543/0,624 | Deleterious/Damaging | 0,0011 | 0,00082 | L12 |
| m.15468C>T | CYB | T241M | 0,890 (possible damaging) | 0,245/0,079 | Neutral/Tolerated | 0,0004 | 0,00043 | L28 |
| m.8551T>C | ATP6 | F9L | 0,976 (probably damaging) | 0,676/0,418 | Deleterious/Damaging | 0,0007 | 0 | L30 |
| m.14582A>G | ND6 | V31A | 0,003 (benign) | 0,245/0,181 | Neutral/Tolerated | 0,0086 | 0,00571 | L40 |
| m.8400T>C | ATP8 | M12T | 0 (benign) | 0,504/0,118 | Neutral/Tolerated | 0,0011 | 0,00052 | L43 |
| m.9444C>T | CO3 | R80W | 0,999 (probably damaging) | 0,875/0,586 | Deleterious/Damaging | 0 | 0 |  |
| m.4639T>C | ND2 | I57T | 0,001 (benign) | 0,297/0,047 | Neutral/Tolerated | 0,0082 | 0,00395 | L50 |
| m.8412T>C | ATP8 | M16T | 0,711 (possible damaging) | 0,677/0,542 | Deleterious/Tolerated | 0 | 0,00039 |  |
| m.15077G>A | CYB | E111K | 0,992 (probably damaging) | 0,684/0,331 | Deleterious/Damaging (low confidence) | 0,0007 | 0,00213 |  |

Phylogenetic analysis clearly illustrates genetic origin of Siberian carriers of pathogenic LHON mutations and confirms that all families are unrelated and belong to different maternal phylogenetic lines. All classic and rare mutations occurred among European haplogroups J, H, V, and U. It is interesting that m.11778G>A were not occurred among haplogroups of branch M. Mutations m.3460G>A and m.14484T>C belong to diverse haplogroups – J, V, H of European origin, M8 and D of Asian origin. Phylogenetic trees are shown in Figures 1, 2 and 3.

**Figure 1, 2, 3.**
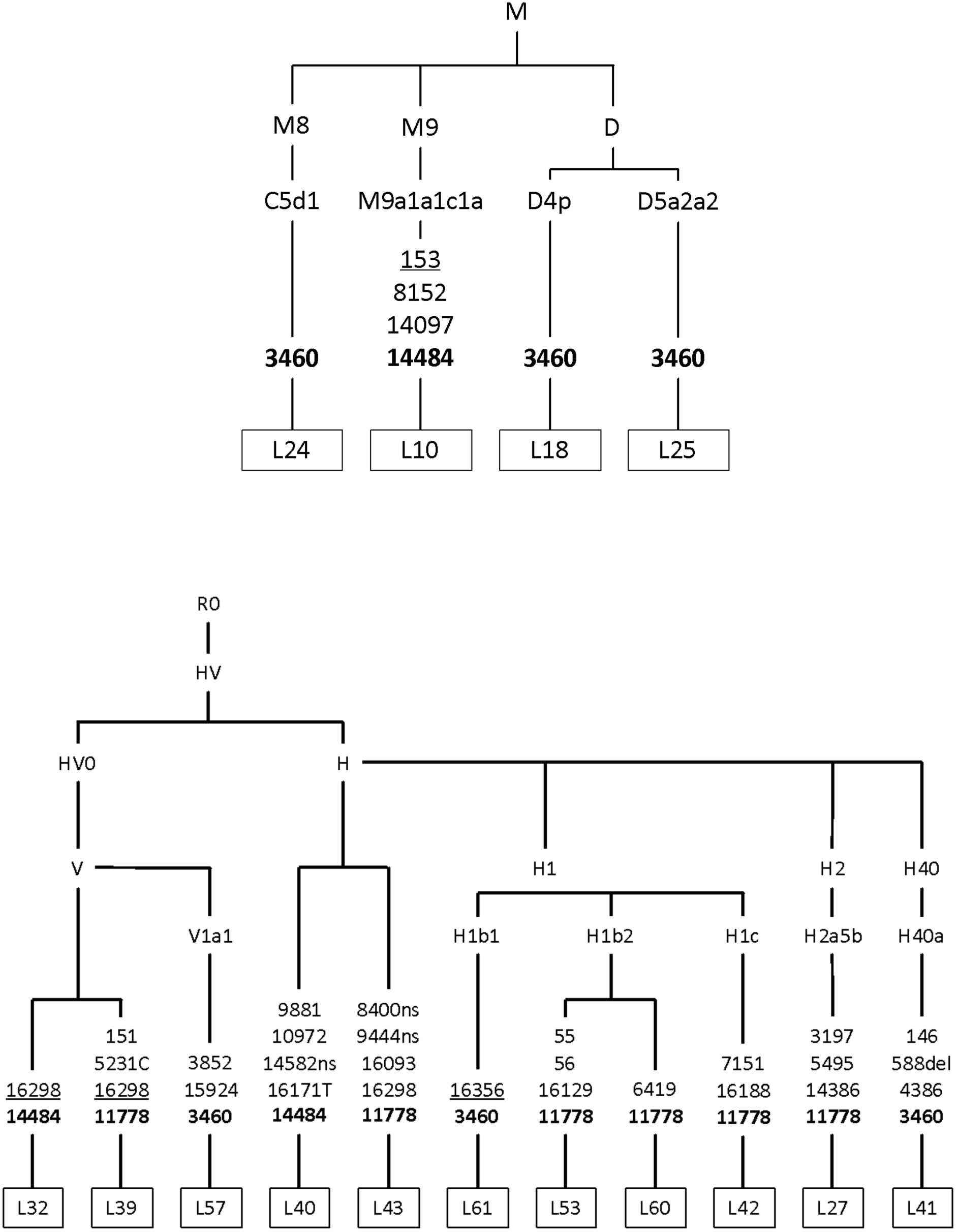
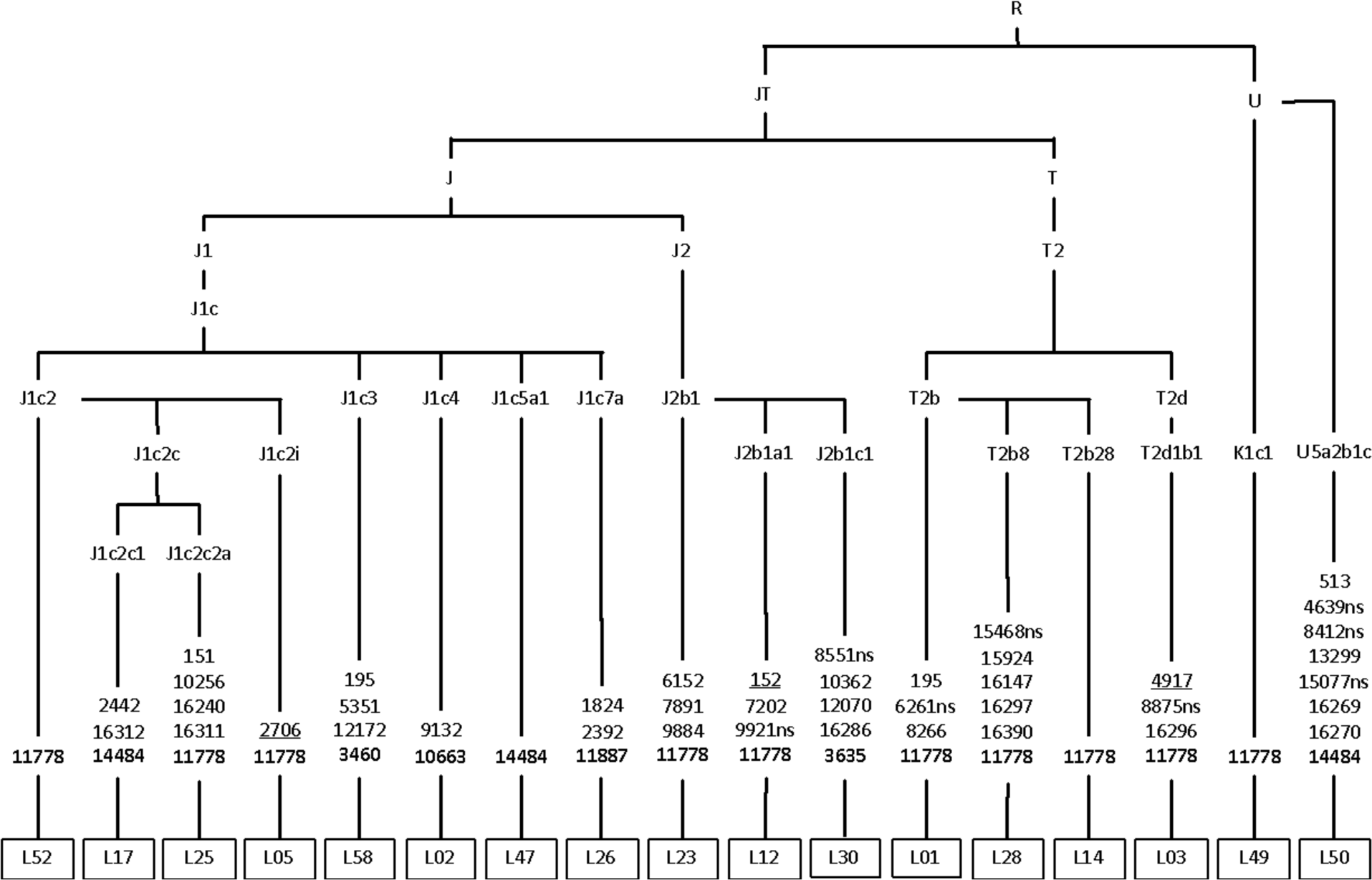
Phylogenetic trees based on complete mtDNA genome sequences of pedigree-probands with pathogenic mutations. The non-synonymous coding-region variants are denoted by “ns” (known pathogenic mutations designated “in bold”), the insertions and deletions are denoted by “ins” and “del,” respectively, underlined mutations are recurrent.

## Discussion

### Primary LHON mutations

In Western Siberia, frequencies of primary mutations are different from frequencies reported for Europe and Asia. The most prevalent m.11778G>A (55%) is less common in Western Siberia than in Europe – 69% (Mackey, Oostra et al. 1996), China and Japan – 90% (Mashima, Yamada et al. 1998, Jiang, Liang et al. 2015). The prevalence of m.3460G>A (24%) is twice as much as in Europe, but m.14484T>C (13,8%) do not deviate from those of other European populations (Mackey, Oostra et al. 1996).

In twelve families with LHON manifestation no one known pathogenic mtDNA mutation was found. However, there are other elaborations in which the clinical diagnosis not confirmed by molecular-genetic analysis (Howell, Oostra et al. 2003, Abu-Amero and Bosley 2006, Aitullina, Baumane et al. 2013, Rosenberg, Norby et al. 2016). Apparently, it is difficult to distinguish the mtDNA mutations-caused LHON from the other forms of optic neuropathy, for example, from the dominant optic neuropathy (DOA), especially when it is the sporadic clinical case without any pattern of inheritance. Compared to LHON, DOA visual loss is detected between ages 4 and 6 in the majority of patients, and 58–84% of patients with DOA report visual impairment by age 11 (Fraser, Biousse et al. 2010). In our 11 cases ages of onset were between 13 – 36 years. We are going to study these cases for the mutation spectrum of common pathogenic genes for DOA in future.

### Penetrance

We suppose that the reasons for low prevalence of LHON disease in Western Siberia are associated with incomplete penetrance and diagnostic difficulties of atypical (e.g. late onset) and combined (e.g. multiple sclerosis) forms of LHON. Patients with LHON could have also been wrongly diagnosed as suffering from toxic amblyopia, tobacco–alcohol amblyopia, or optic neuritis (Rosenberg, Norby et al. 2016). On the other hand, the problem is that patients do not now their family history. Molecular testing for LHON is not routinely performed in patients with optic atrophy in Russian Federation. Identification and registration of unaffected carriers plays an important role for prevention of disease manifestation. For example, there is strong evidence now that smoking is associated with an increased risk of visual failure among LHON carriers – 93% penetrance of vision loss in male smokers versus a 66% penetrance in male non-smokers (Kirkman, Yu-Wai-Man et al. 2009). Our observations highlight the importance of molecular genetic examination for unaffected carriers. Since the proportion of sporadic cases is high, 39% according to our data, 40% according to published data (Meyerson, Van Stavern et al. 2015), the presence of pathogenic mutations should be tested not only for probands but for all relatives on the maternal line.

### Potentially pathogenic mutation m.4659G>A

We found out m.4659G>A in one subject without any known primary mutations (L51). This sequence change is located at codon 120 in the functional domain of the CO1 gene and changes an alanine, a hydrophobic amino acid, into threonine - a neutral amino acid. Mutation m.4659G>A has been reported as associated with LHON in an Australian pedigree that also had heteroplasmic mutation m.14484T>C and m.5460G>A (Mackey and Howell 1992). This family had 10 maternally related descendants, five of whom with vision loss. Unfortunately, our patient does not know his family history and we couldn’t confirm maternal inheritance for this mutation. However, m.4659G>A has very low frequency in the general population (0,0011) and high probability to be pathogenic according to data of different prediction tools (Table 3).

### Additional non-synonymous mutations revealed in LHON cases

Phenomenon of co-existence of two pathogenic mutations in one family has already been described. In the first case, m.14484T>C, m.4659G>A and m.5460G>A in an Australian LHON pedigree, described above (Mackey and Howell 1992). In the second case, a family harboring two primary LHON mutations m.11778G>A and m.14484T>C, and both mutations had synergistic pathogenic effect on protein function, and a higher degree of heteroplasmy of the m.14484T>C correlated with an earlier age at onset (Catarino, Ahting et al. 2016). And the third example is a unique double-mutant ND4L with two concurrent mutations m.10609T>C and m.10663T>C in an Arab pedigree from Kuwait (Behbehani, Melhem et al. 2014).

We reported mutations m.9444C>T, m.6261G>A, m.9921G>A, m.8551T>C, m.8412T>C, m.15077G>A, which could be potentially pathogenic because of low frequency in the general population and high probability to be pathogenic according to data from different prediction tools. Two of them m.6261G>A and m.15077G>A have already been reported in subjects with optic neuropathy and maternally inherited isolated deafness respectively. However, we suppose that additional non-synonymous mutations could either have a synergistically pathogenic or a protective effect. To demonstrate the full significance of novel mutations respiratory chain assay should be performed. For example, as in the study by Gutierrez Cortes, Pertuiset et al. 2012, where cybrids with m.15077G>A showed normal activities for mitochondrial electron chain enzymatic complexes.

### Haplogroup analysis

Western Siberia region is represented predominantly by European haplogroups and includes aboriginal Asian populations. Asian haplogroups have been found in Altaian (L18) and Tuvinian (L24) as in Russian families (L10, L25).

Rare mutations m.10663T>C and m.3635G>A were found in Russian families from Kazakhstan (the first) and the Novosibirsk region (the second) associated with the European haplogroups J1c4 and J2b1c1, respectively (Brown et al. 2001, Brown et al. 2002). Mutation m.10663T>C was also reported in the background of the haplogroups J1c2c, L2a1, L3’4, L3f1b (Abu-Amero and Bosley 2006, Achilli, Iommarini et al. 2012, Behbehani, Melhem et al. 2014, Al-Kharashi, Al-Kharashi et al. 2016) and mutation m.3635G>A - R11a, D4g2b, M7b4, F1a, B5b, M7b (Yang, Zhu et al. 2009, Bi, Zhang et al. 2012). The presence of the same pathogenic mutations on the background of various mitochondrial haplogroups confirms that pathogenic LHON mutations arise de novo, independently of different mtDNA haplogroups and ethnic backgrounds. It is known that the clinical impact of mDNA mutations may be modulated by mitochondrial haplogroups. For example, Hudson et al. performed a multicenter study of 3,613 subjects from 159 different families and showed that the risk of visual failure is greater when m.11778G>A or m.14484T>C mutations are present in specific subgroups of haplogroup J, the m.3460G>A mutation is present in haplogroup K, and the risk of visual failure is significantly lower when m.11778G>A occurs in haplogroup H (Hudson, Carelli et al. 2007) Romero et al. supposed that haplogroup D has a protective effect in carriers of LHON mutations. His hypothesis was based on the fact that there was a markedly decreased frequency of haplogroup D in Chilean subjects with LHON, as haplogroup D is one of the most common in the Chilean population (Romero, Fernandez et al. 2014). Also, other experimental research serves as proof that cybrids and fibroblasts bearing LHON mutations have different response to neurotoxic agents depending on haplogroup background (Ghelli, Porcelli et al. 2009).

Association of mtDNA variation in different diseases could be a result of founder events. It is suggested, that at the end of the last glaciation phylogenetically more ancient mutations could have provided their carriers with the adaptive advantages upon the development of Central and Northern Europe, and positive selection could contribute to the fixation of weakly pathogenic LHON mutations, appearing at specific genetic background (Volod’ko, L’vova M et al. 2006).

## Conclusion

These data may be insufficient for evaluating the prevalence of LHON. However, the project is still ongoing, and all the data collected might be used to develop a LHON registry in Russian Federation.

